# Robust expansion of phylogeny for fast-growing genome sequence data

**DOI:** 10.1101/2021.12.30.474610

**Authors:** Yongtao Ye, Marcus H. Shum, Joseph L. Tsui, Guangchuang Yu, David K. Smith, Huachen Zhu, Joseph T. Wu, Yi Guan, Tommy T. Lam

## Abstract

Massive sequencing of SARS-CoV-2 genomes has led to a great demand for adding new samples to a reference phylogeny instead of building the tree from scratch. To address such challenge, we proposed an algorithm ‘TIPars’ by integrating parsimony analysis with pre-computed ancestral sequences. Compared to four state-of-the-art methods on four benchmark datasets (SARS-CoV-2, Influenza virus, Newcastle disease virus and 16S rRNA genes), TIPars achieved the best performance in most tests. It took only 21 seconds to insert 100 SARS-CoV-2 genomes to a 100k-taxa reference tree using near 1.4 gigabytes of memory. Its efficient and accurate phylogenetic placements and incrementation for phylogenies with highly similar and divergent sequences suggest that it will be useful in a wide range of studies including pathogen molecular epidemiology, microbiome diversity and systematics.

## Introduction

Next generation sequencing (NGS) technologies enable large-scale exploration of diversity and monitoring temporal evolution of organisms, which often involve generating and analyzing large numbers of sequences from new organisms on an ongoing basis. For instance, over 5 million of SARS-CoV-2 genomes have been sequenced within two years of the pandemic (Shu & McCauley, 2017), largely facilitating transmission tracking and disease control. Conventional methods of phylogeny inference from scratch such as those implemented in IQ-TREE2 (Minh et al., 2020) and FastTree2 (Price, Dehal, & Arkin, 2010) could hardly cope with such rapidly growing huge sequence datasets. Therefore, determining the evolutionary position of new sequences as they become available by placing or inserting them into the reference tree becomes a more efficient alternative. Such ‘phylogenetic placement’ has been useful for taxonomic classification, while accumulative addition of sequences (incrementing the phylogeny as a result) allow efficient update of the growing phylogeny in a global context.

Previously published methods such as PhyClass (Filipski, Tamura, Billing-Ross, Murillo, & Kumar, 2015), EPA-ng (Barbera et al., 2019) and pplacer (Matsen, Kodner, & Armbrust, 2010) utilize minimum evolution or maximum likelihood criteria to infer the evolutionary position of the query sequence and place it directly onto a pre-built phylogeny. These algorithms require relatively large computer memory or long runtime which makes massive sequence insertion difficult. Recently, in respect of tracking the diversity of the large amount of SARS-CoV-2 virus genomes, UShER (Yatish Turakhia et al., 2021) was developed to tackle this problem by calculating the ‘branch parsimony score’ to search for positions of taxa placement only based on sequence mutations to a particular reference. It is extremely fast as compared to the other existing programs. Although the performance of UShER on the SARS-CoV-2 genomes is promising, the placement performance for genome sequences with greater divergence is not well studied.

We hereby introduce a new approach TIPars, which inserts sequences into a reference phylogeny based on parsimony criterion with the aids of a full multiple sequence alignment of taxa and pre-computed ancestral sequences. The ancestral sequences are useful and efficient in assisting the search of the best placed position because these ancestral sequences often contain rich information in the evolution context of a phylogenetic tree (Loytynoja, Vilella, & Goldman, 2012). Recent ancestral sequence reconstruction methods such as PastML (Ishikawa, Zhukova, Iwasaki, & Gascuel, 2019) and RASP4 (Y. Yu, Blair, & He, 2020) have improved speed and accuracy to become feasible in the huge SARS-CoV-2 phylogeny. TIPars searches the position for insertion by calculating the triplet-based minimal substitution score for the query sequence on all branches (Fig. 1A). To compare the performances of different phylogenetic placement/insertion methods including TIPars, UShER, EPA-ng, IQ-TREE2 and PAGAN2 (Loytynoja et al., 2012), we applied them on four benchmark datasets (SARS-CoV-2, Influenza virus, Newcastle disease virus and 16S rRNA genes). The first test is single taxon placement. We pruned one taxon from a given phylogenetic tree and applied the methods to place it back. The second is multiple taxa insertion in which a set of taxa was removed and sequentially inserted back. We compared the topology and log likelihood for the trees before pruning and after reinsertion. Our evaluation tests aimed to assess the robustness of the methods on both highly similar sequences and divergent sequences, and whether the phylogenetic tree could be efficiently updated with new sequences that are continuously generated.

**Fig. 1.**
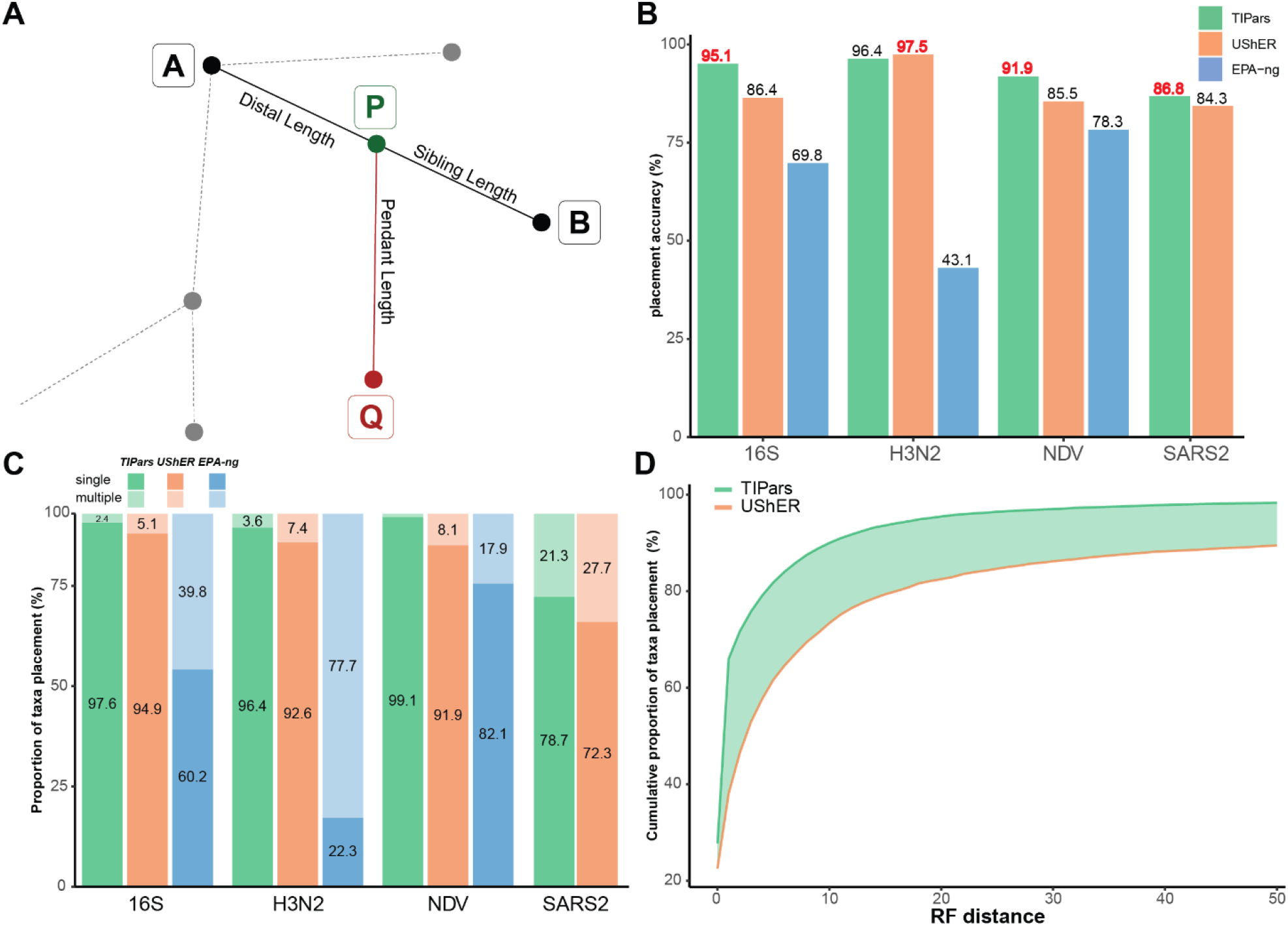
Illustration of phylogenetic placement and single taxon placement performance. (**A**) Illustration of the placement for a query sequence. “Q” indicates the query sequence, “A” and “B” represent the existing nodes in the reference tree. “P” represents the parental node of “Q” generated by TIPars. Minimum substitution score is calculated based on the triplet formed by A-B-Q. (**B**) Bar charts represent the accuracy of single taxon placement on 16S, H3N2, NDV and SARS2-100k datasets using TIPars, UShER and EPA-ng respectively. Accuracy is indicated on top of each bar and the highest accuracy in each dataset is highlighted in red. (**C**) Stacked bar charts show the proportion of single and multiple taxon placement result for TIPars (Green), UShER (Orange) and EPA-ng (Blue) on the 16S, H3N2, NDV and SARS2-100k datasets. Proportion with > 1% is indicated within the bar. (**D**) Cumulative proportion of single taxa placement on the SARS2-660k dataset with different RF distance cutoff. Highlighted area represents the difference between TIPars and UShER.

## Results

### Computational performance of TIPars and other methods

A number of approaches have been proposed for phylogenetic placement or insertion, but dealing with the vast number of SARS-CoV-2 genome sequences has rendered most of these methods impractical or computationally prohibitive. Based on a reference SARS-CoV-2 phylogenetic tree (SARS2-100k) generated from 96,020 unmasked SARS-CoV-2 sequences of high quality (details in Methodology), we evaluated our proposed program TIPars with UShER, EPA-ng, IQ-TREE2 and PAGAN2 by sequentially inserting 100 new sequence samples. Only TIPars and UShER were practicable in terms of running time and memory usage. PAGAN2 were not able to complete the insertion within 96 hours and hence no data was available. Although IQ-TREE2 used a lower peak memory than EPA-ng, the running time was the highest among all programs. In contrast, EPA-ng achieved a faster running time than IQ-TREE2 but the peak memory usage was around 1 terabyte (TB) which would not be practicable for general users. As for TIPars, it took only 21 seconds (excluding the input loading time) on a 64-cores server and required about 1.4 gigabytes (GB) peak memory usage (Table 1). Another computational performance comparison on smaller dataset with 800 bacterial 16S rRNA sequences (16S) can be checked in table S1 in which PAGAN2 was runnable. Overall, in the SARS2-100k phylogenetic tree, TIPars ran 10-300 folds faster than EPA-ng and PAGAN2 with 98.5% to 99.9% less memory used, an efficiency that is comparable to that of the leading program UShER.

**Table 1.**
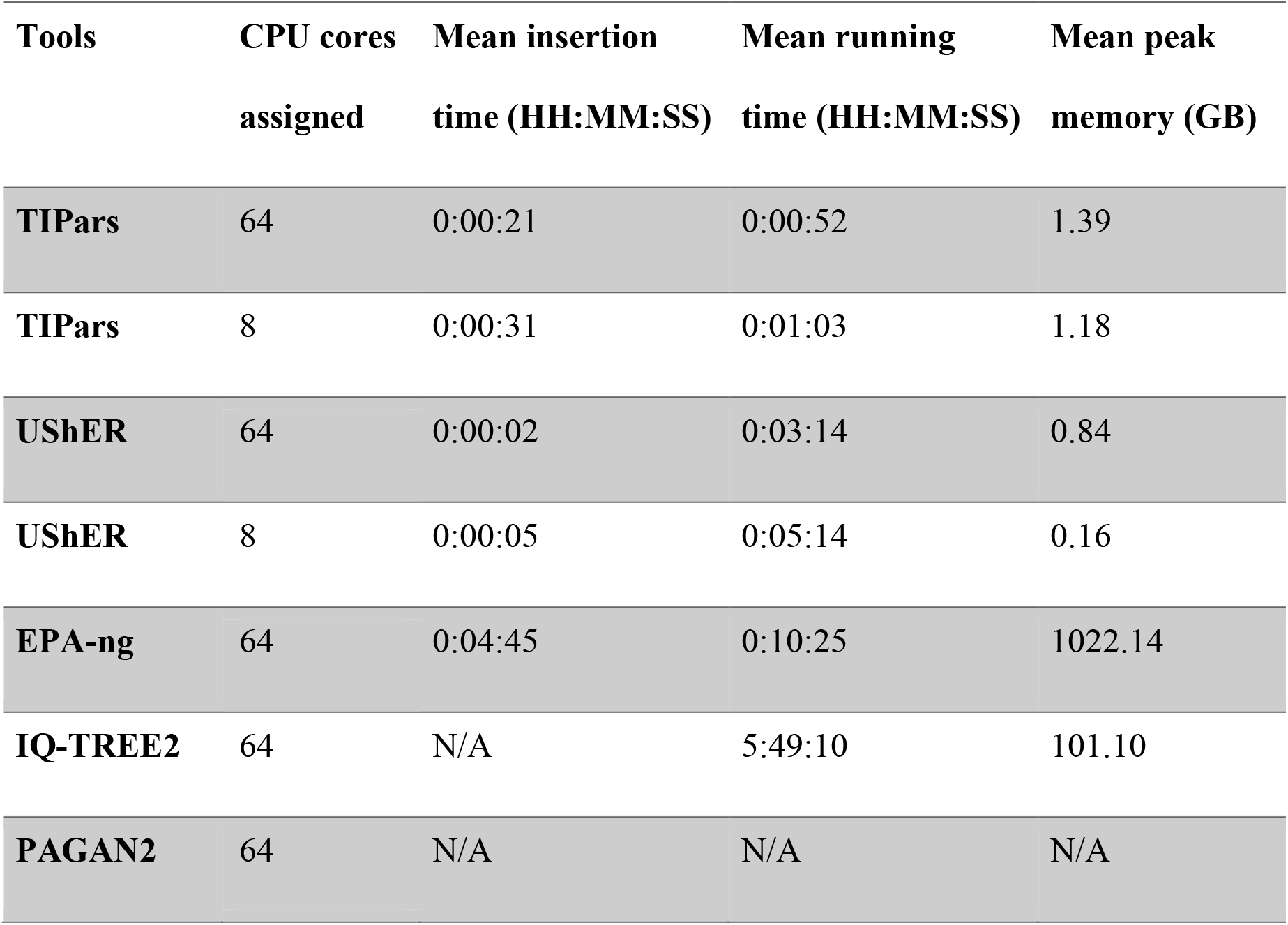
Average running time and memory used through 10 repeated runs of inserting/placing 100 genome samples into SARS2-100k reference tree. Tests were running on a server of 64 Intel Xeon Gold 6242 CPU cores and 1500 GB RAM. We also compared TIPars with UShER on a general computer with 8 CPU cores. TIPars ran with a JAVA setting of Xmx1G. The running time of UShER contains its necessary computation of ‘mutation-annotated tree’. PAGAN2 was not runnable for this dataset. N/A indicates that data are not applicable.

| **Tools** | **CPU cores**<br>**assigned** | **Mean insertion**<br>**time (HH:MM:SS)** | **Mean running**<br>**time (HH:MM:SS)** | **Mean peak**<br>**memory (GB)** |
| --- | --- | --- | --- | --- |
| **TIPars** | 64 | 0:00:21 | 0:00:52 | 1.39 |
| **TIPars** | 8 | 0:00:31 | 0:01:03 | 1.18 |
| **UShER** | 64 | 0:00:02 | 0:03:14 | 0.84 |
| **UShER** | 8 | 0:00:05 | 0:05:14 | 0.16 |
| **EPA-ng** | 64 | 0:04:45 | 0:10:25 | 1022.14 |
| **IQ-TREE2** | 64 | N/A | 5:49:10 | 101.10 |
| **PAGAN2** | 64 | N/A | N/A | N/A |

### Single taxon placement

Adding a single sequence sample (query) into a reference tree is useful to obtain the phylogenetic placement of the new data, and can be the basic step for expanding the phylogeny with new sequences. We tested TIPars, UShER and EPA-ng on four datasets, including the SARS-CoV-2 genomes (SARS2-100k), 16S ribosomal RNA genes (16S), hemagglutinin genes of human seasonal influenza A viruses (H3N2), and Newcastle disease virus genomes (NDV) where the average pairwise genetic distances (substitutions per site) of SARS2-100k and H3N2 are less than 0.04 (similar sequences) while those of 16S and NDV are greater than 0.12 (divergent sequences) (details in Methodology; table S2). For the SARS2-100k dataset, EPA-ng was not applied due to impractically large memory requirement and long runtime.

Based on the postorder traversal, between every 10 taxa we selected one sequence from the SARS2-100k sequence alignment resulting in 9,602 sequences, i.e., 10% of the total taxa in the tree. These selected sequences were individually removed from the reference tree and multiple sequence alignment (MSA) one at a time and used as the query sample for single taxon placement. In datasets of 16S, H3N2 and NDV, all taxa were removed individually and used for the placement test.

To evaluate the accuracy of each single taxon placement, we calculated the Robinson-Foulds (RF) distance (Robinson & Foulds, 1981) between the reference tree before the taxon removal and the resulting tree after the placement using corresponding programs. An RF distance measures the topological clustering difference between two trees. A zero RF distance indicates that the two trees are topologically identical, and hence the single taxon placement position is exactly the same as the original position, i.e. a true positive.

With the aid of ancestral information and MSA of full sequences, TIPars performed accurately on phylogenies made of highly similar (SARS2-100k and H3N2) and divergent (16S and NDV) sequences (Fig. 1B). However, a drop in accuracy on more divergent sequences was observed from UShER, perhaps because UShER was only based on the sequence mutations to a particular reference sequence as input, which may lose the insertion information (Yatish Turakhia et al., 2021). In addition, we noted that due to the massive sequencing of SARS-CoV-2 by different research groups, sequencing quality varies and ambiguity bases often occur in the consensus genome sequence data, which could affect the placement accuracy. To account for ambiguity data in sequencing, we used a specific substitution scoring table based on the IUPAC nucleotide ambiguity codes (table S3) for the taxon placement and insertion process (details in Methodology), which achieved a robust performance in sequences of different qualities.

Notably, when searching through the whole phylogeny for the best position to place a taxon, there may be cases where multiple branches achieve equal minimum substitution scores, and thus the placement will be uncertain. As demonstrated in Fig. 1C, TIPars produced the least number of multiple ambiguously optimal placements in all testing datasets. For example, TIPars generated 23% fewer multiple placements than UShER in the SARS2-100k dataset.

A possible reason for the relatively poor performance of EPA-ng could be that RF distance may not be a reliable metric to compare binary trees derived from the phylogeny with polytomy because there is a very skewed distribution of RF distance when comparing two random binary trees (Bryant & Steel, 2009; Lin, Rajan, & Moret, 2012; Moon & Eulenstein, 2019). It is notable that EPA-ng only processes binary trees. To address this issue, a relaxed criterion for true positive was applied based on whether there are common sister taxa for the removed and re-placed single taxon, as previously used (Yatish Turakhia et al., 2021). With the adjusted true positive measurement, TIPars achieved the highest accuracy in all datasets (fig. S1). While the accuracy of EPA-ng was substantially improved, it was still the lowest among the three tested programs.

To assess the practicability for extremely large phylogenies, we applied TIPars and UShER in single taxon placement test over the global SARS-CoV-2 phylogenetic tree with 659,885 masked genome sequences (SARS2-660k) downloaded from the Global Initiative on Sharing All Influenza Database (GISAID) (Shu & McCauley, 2017) on the 6th September 2021. A total of 65,989 sequences (10% of the total taxa in the tree) were removed and re-inserted individually. Cumulative proportion of single taxon placement result with different RF distance cutoff was shown in Fig. 1D. TIPars produced trees with significantly higher topological similarity to the reference tree with a median RF distance of 0.5 and mean of 5.8 (99% confidence interval (CI) = [5.5-6.1]) as compared to UShER (median RF distance is 3.0 and mean is 31.2 (99% CI = [30.03-2.4])) at 99% significance level (p-value < 10^-10^).

### Multiple taxa insertion

Multiple taxa insertion was an alternative method in determining the phylogenetic position of new sequences over conventional complete phylogeny construction from scratch. TIPars and other three programs (IQ-TREE2, PAGAN2 and UShER) were applied on the four datasets to conduct a comprehensive evaluation of performance.

In the SARS2-100k dataset, we performed multiple taxa insertion for 100 sets of 10^2^ and 10^3^ randomly selected sequences (an example is shown in Fig. 2A) (random100 and random1000) and 100 sets of 10^2^ and 10^3^ successively selected sequences (i.e., a set of successive taxa following the tree postorder traversal; an example is shown in Fig. 2B) (successive100 and successive1000). In the 16S, H3N2 and NDV datasets, 100 sets of 50 sequences were randomly selected. The selected sequences are pruned from the corresponding reference tree and become multiple taxa to be reinserted for each testing set.

**Fig. 2.**
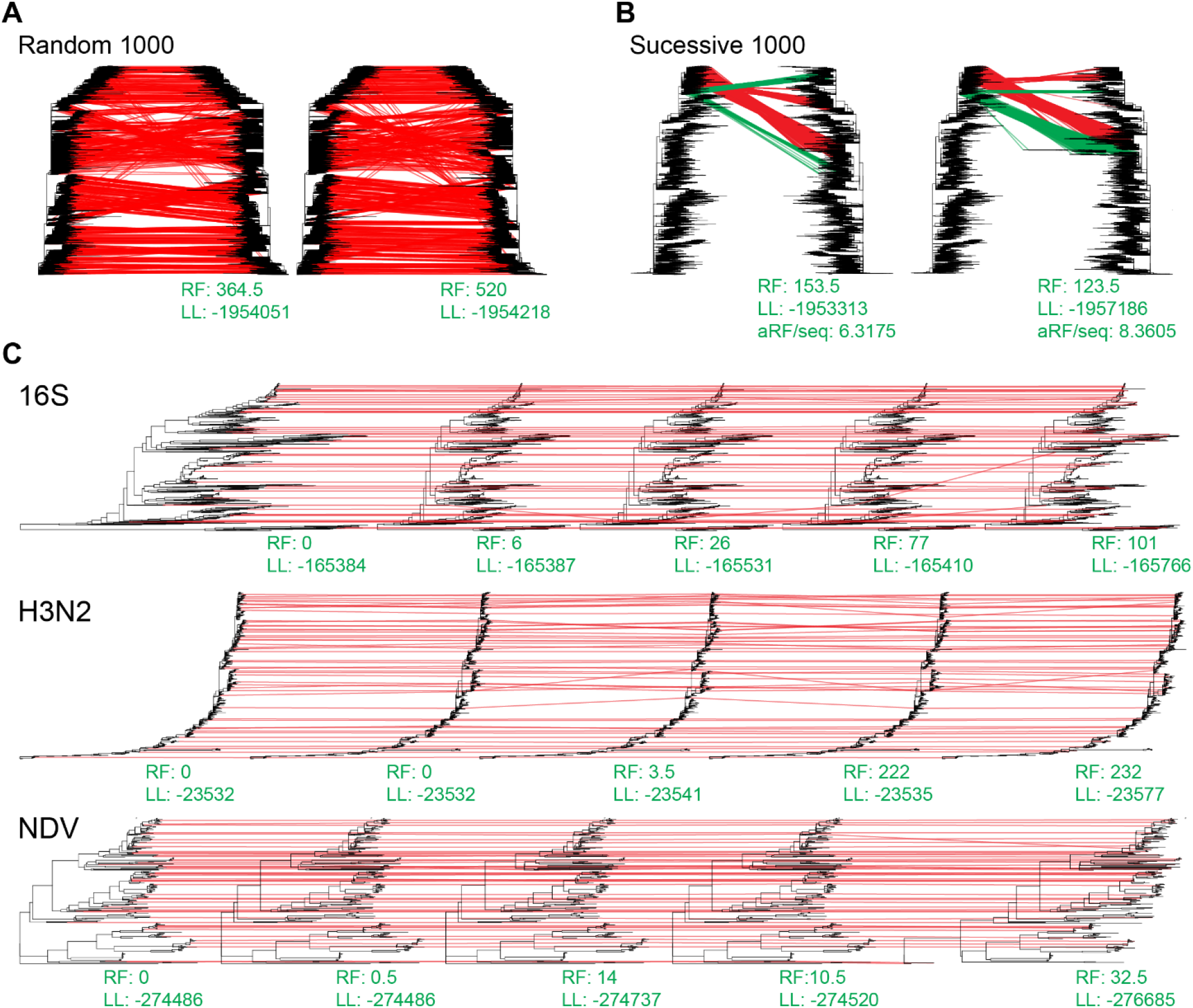
Taxa insertion visualization. (**A**) A demonstration of TIPars resulting tree (Left) and UShER resulting tree (Right) paired with the reference SARS2-100k reference tree (Left tree in both figures) for the insertion of randomly selected 1000 taxa sequences. Red lines link the corresponding positions of inserted taxa between reference and resulting tree. (**B**) A demonstration of TIPars resulting tree (Left) and UShER resulting tree (Right) paired with the reference SARS2-100k reference tree (Left tree in both figures) for the insertion of successively selected 1000 taxa sequences. Green lines indicate different taxa insertion positions between TIPars and UShER. Averaged RF-distance per sequence (aRF/seq) comparing to the reference tree is shown at the bottom. (**C**) Demonstrations of the resulting trees for randomly selected 50 taxa in NDV, 16S (Midpoint rooted) and H3N2 datasets. From the left to the right are trees of reference, TIPars, UShER, IQ-TREE2 and PAGAN2. RF distance (RF) compared to the reference tree and the Gamma20 log-likelihood (LL) are shown at the bottom of each resulting tree.

RF distance and tree log-likelihood (LL) were used to evaluate the performance of the multiple sequence insertion. To evaluate the topology accuracy, the resulting tree produced by the four programs were compared to the original reference tree (leaf taxa unpruned) to obtain the RF distance. At the same time, Gamma20 log-likelihoods of the reference tree and the resulting tree after optimizing the branch length were also computed using FastTree2 (double-precision version) and their differences were used for evaluation.

For the random100 and random1000 datasets, only analyses using TIPars and UShER were able to complete within a reasonable computation time, hence no result from IQ-TREE2 and PAGAN2 was present. The resulting trees from multiple taxa insertion using TIPars had a significantly smaller RF distance than those generated using UShER (Fig. 3A). In addition, the log-likelihood of the resulting trees from TIPars was significantly higher than that of UShER (Fig. 3B). Moreover, TIPars resulting trees tended to be very close to the reference tree with smaller log-likelihood differences (fig. S2, A and B). A demonstration of the taxa-insertion was illustrated in Fig. 2A by adding 1000 samples. We observed there were more crossing lines from reference tree to UShER resulting tree indicating more misplaced insertions.

**Fig. 3.**
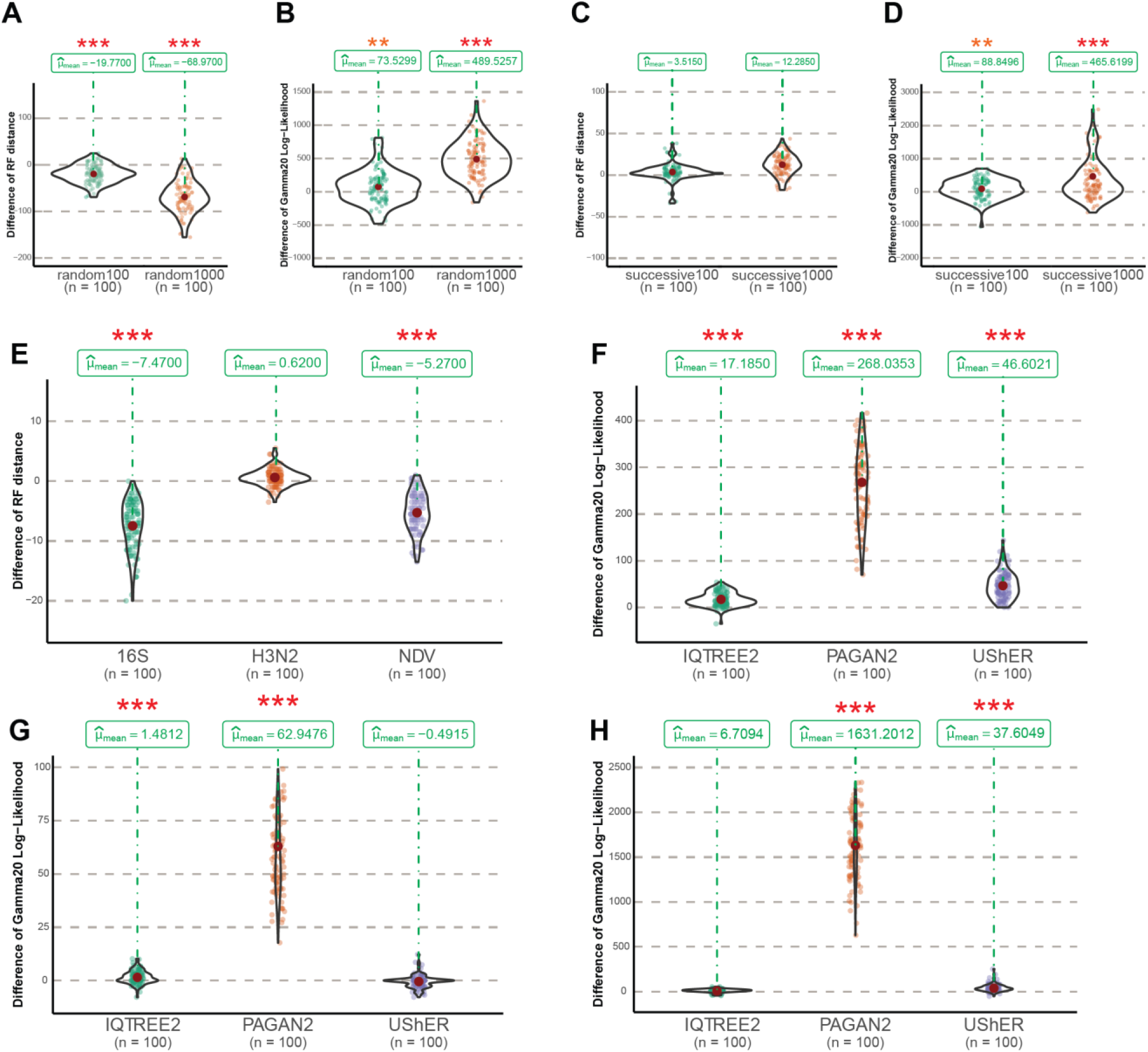
Multiple sequences insertion performance. (**A-D**) Violin graphs show the distribution of paired differences of the RF distance and the Gamma20 log-likelihood between the optimized resulting trees generated by TIPars and UShER (TIPars - UShER) for the random 100, 1000 and successive 100 and 1000 multiple sequences insertions. (**E**) Distribution of the paired difference of the RF distance between the optimized resulting trees generated by TIPars and UShER (TIPars - UShER) on 16S, H3N2 and NDV random 50 multiple sequences insertions. (**F-H**) Distribution of the paired difference of the Gamma20 log-likelihood between the optimized resulting trees generated by TIPars and the three other programs (TIPars - Others) on 16S (**F**), H3N2 (**G**) and NDV (**H**) random 50 multiple sequences insertions. P-value for the right-sided paired t-test is indicated by the asterisk on top of each violin diagram, where p<0.05 is indicated by one pink asterisk (*), p<0.01 by two orange asterisks (**) and p<0.001 by three red asterisks (***).

As to 16S, H3N2 and NDV datasets, TIPars mostly outperformed IQ-TREE2, PAGAN2 and UShER with a significantly lower RF distance and a higher log-likelihood of resulting trees (Fig. 3, E to H; fig. S3). In the H3N2 dataset, there was no significant tree log-likelihood difference between TIPars and UShER (Fig. 3G), and in NDV dataset, TIPars performed better than IQ-TREE2 with higher mean log-likelihood but without statistical significance (Fig. 3H). The demonstrations of the taxa-insertion result were visualized in Fig. 2C where UShER, IQ-TREE2 and PAGAN2 were less accurate than TIPars.

For the successive100 and successive1000 datasets, TIPars resulting trees had a significantly larger RF distance than those of UShER (Fig. 3C). However, the log-likelihood of the TIPars resulting trees was significantly higher than that of UShER (Fig. 3D; fig. S2, C and D). By comparing the trees generated from TIPars and UShER (Fig. 2B), the difference is that TIPars inserted some of query taxa (green lines in Fig. 2B; successive taxa pruned from the reference tree) into two subtrees where one of them (the one containing over half the queries) had the same topology as the one in the reference tree. Whereas UShER inserted those queries mostly within a monophyletic clade but it was different from the reference tree. As a result, UShER retained the local topology (better RF distance) (Lin et al., 2012; Smith, 2021) but missed the global topology (worse log-likelihood). Through a RF distance comparison specifically to each query taxon instead of all query taxa, we found that the RF distance resulted from UShER was not significantly higher than that of TIPars (table S4).

On the other hand, we may suppose that in the situation of random100 and random1000 tests, RF distance would be a suitable metric for comparing the performance of taxa insertions as they are similar to the case of single taxon placements, where most removed taxa are within different monophyletic clades due to randomness (Bryant & Steel, 2009).

To make the log-likelihood of the resulting trees comparable, we applied FastTree2 to reoptimize the branch lengths with fixed topology (Price et al., 2010). However, compared to the efficiency of taxa insertion (Table 1), the re-optimization is time-consuming. For example, the optimization for a SARS2-100k tree took 10 to 12 hours and required around 125 GB memory (table S5). Therefore, we also computed the log-likelihoods with fixed branch lengths (FLL) using IQ-TREE2, and TIPars still outperformed UShER significantly (fig. S4) by achieving a higher log-likelihood in the resulting tree output directly from the program.

### Inserting novel sequences

To verify practicability of TIPars in adding novel sequences into a given phylogeny, we further performed an experiment to insert novel real-world SARS-CoV-2 samples into the SARS2-100k reference tree. We randomly selected SARS-CoV-2 samples from GISAID which were not included in the SARS2-100k dataset. Twenty sets of 100, 1000, 5000 and 10000 genome sequences were generated as the queries for taxa insertion using TIPars and UShER.

Log-likelihoods of the resulting trees from each program were calculated and their pairwise differences between TIPars and UShER were used to evaluate the performance. RF distance was not a suitable metric in this experiment as a comparable reference tree was not available. TIPars provided a resulting tree with a significantly better log-likelihood than UShER in all situations (p-values <0.05; Fig. 4A).

**Fig. 4.**
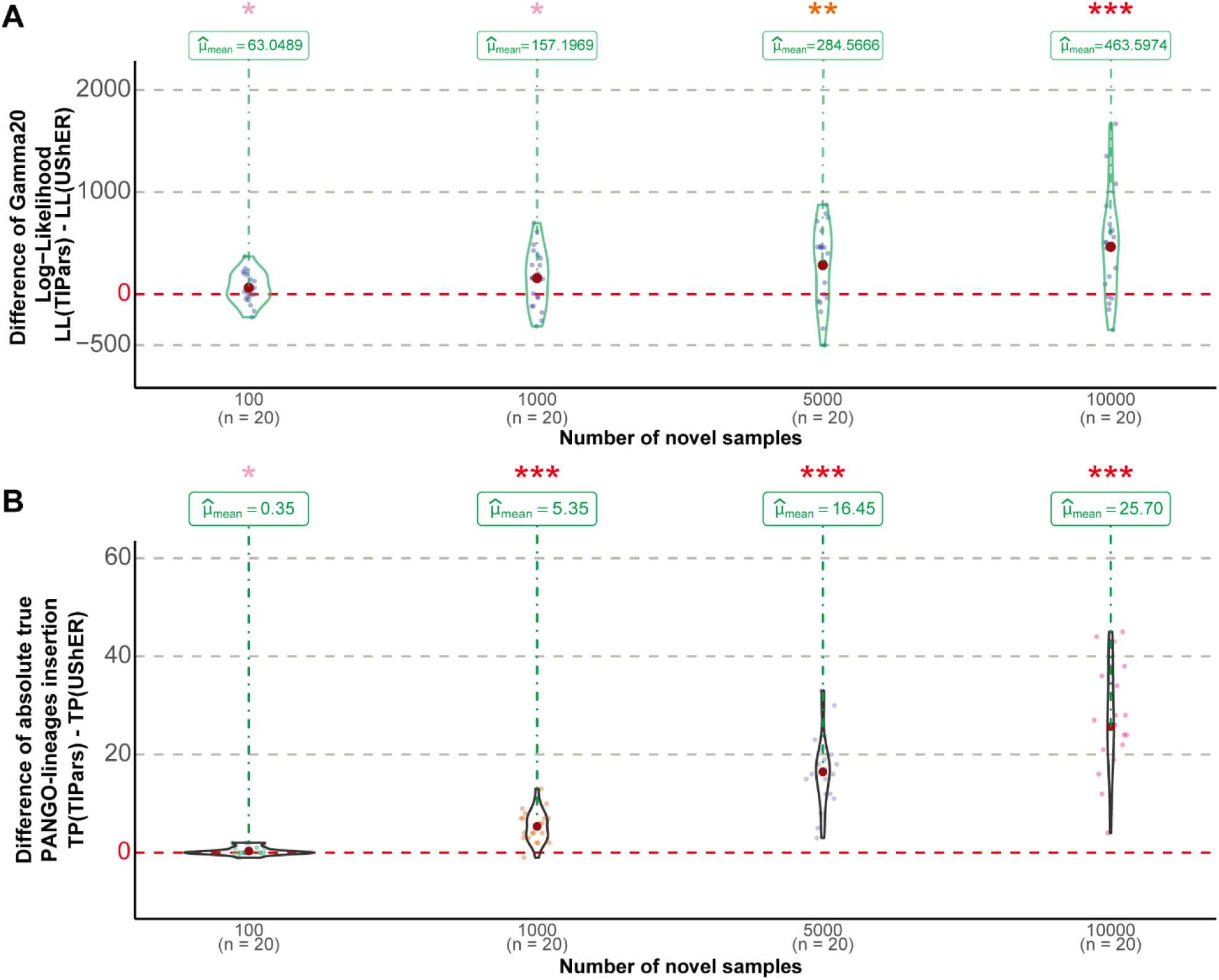
Performance of inserting actual novel sequences. (**A, B**) Violin graph represents the distribution of the paired differences between the Gamma20 log-likelihood (LL) **(A)** and the absolute number of true PANGO-lineages insertion (TP) **(B)** of TIPars over UShER. p-value for the right-sided paired t-test was indicated by the asterisk on top of each violin diagram, where p<0.05 indicated by one pink asterisk (*), p<0.01 by two orange asterisks (**) and p<0.001 by three red asterisks (***).

In addition to tree log-likelihood, we also compared the PANGO lineages (PANGOlins) assignment of the added samples (Rambaut et al., 2020) to validate the accuracy. Only PANGOlins that existed in the reference tree were considered. We assigned each newly inserted sequence with the lineage name of the subtree under the parental node of the inserted position. The subtree was annotated by its descendant reference taxa if all of them were monophyletic (McBroome et al., 2021). A true positive was defined as when the assigned lineage of a query sequence was identical to its original PANGOlins label. In case of queries within unannotated subtrees, we ignored them in the calculation. TIPars outperformed UShER by achieving higher true positive samples on the 100, 1000, 5000 and 10000 insertion datasets with an average of 92% PANGOlins accuracy. The superiority of TIPars was statistically significant under a right-tailed paired t-test (p-values < 0.001) on the 1000, 5000 and 10000 datasets (Fig. 4B and table S6).

## Discussion

TIPars showed promising taxa placement and insertion accuracy in the phylogenies with homogenous (H3N2 and SARS2-100k) and divergent (16S and NDV) sequences, and in extremely large phylogeny (SARS2-660k) with reasonable runtime and memory usage. Although UShER has a lower accuracy in the divergent sequence datasets (16S and NDV), it ran faster than TIPars (Table 1).

Reconstruction of ancestral sequences are associated with all taxa across the phylogenetic tree, which could be done using maximum likelihood statistical models or other advanced techniques (Ishikawa et al., 2019; Kosakovsky Pond et al., 2020; Pupko, Pe’er, Shamir, & Graur, 2000; Y. Yu et al., 2020). So ancestral sequences may reveal more accurate (especially intermediate) evolutionary information than the consensus mutation lists along each individual lineage as UShER does. The evolutionary information can be used to distinguish insertion, deletion and substitution events in the searching of taxon placement (Löytynoja & Goldman, 2005), which may help TIPars to be robust on more divergent phylogenies (Loytynoja et al., 2012). Overall, compared to existing phylogenetic placement programs, TIPars is a robust method for a variety of datasets with densely sampled and highly similar sequences of a single species which are common in tracking pathogen epidemiology and transmission, as well as the sequences with greater intraspecific divergence such as the genome datasets at genus, families or higher taxonomic levels for systematics studies.

Although we showed that TIPars resulting trees with higher tree log-likelihood compared to other programs, a general limitation of the phylogenetic placement method is that errors from incorrect placements accumulate as multiple sequences are inserted sequentially. In order to minimize the error due to large numbers of sequence insertions, it is suggested to conduct tree refinements on not only branch length but also tree topology using different techniques such as nearest-neighbor interchanges (NNIs) and subtree-pruning-regrafting (SPRs) (Price et al., 2010). Furthermore, starting such optimization process with an initial tree of higher log-likelihood may achieve a final tree with better log-likelihood using certain of time (Price et al., 2010). As demonstrated in table S7, for the resulting trees of equal RF distance from both TIPars and UShER (n=28), the branch length optimized trees for TIPars had higher (n=14) or equal (n=12) tree log-likelihoods than the ones resulted from UShER.

TIPars could facilitate the future development of sequence analysis methods that make use of the phylogenetic placement information. For instance, genome assembly of NGS read data from the metagenome can use phylogenetic positions of the short-read sequences to distinguish between related microbial strains or lineages. With the aid of TIPars, NGS sequences could be inserted to the branches of specific strains or lineages in a reference phylogeny. This can be used in calculating the proportion of strains in mixed infection even when one of the strains is at low abundance in which *de novo* assembly may generate incomplete contigs.

Since the start of the COVID-19 pandemic, over 5 million SARS-CoV-2 genome sequences have been made publicly available (Shu & McCauley, 2017). With the reduction in cost, the rate of genome sequencing is expected to skyrocket in the future. By providing rapid and memory efficient taxa insertions at high accuracies, TIPars may improve real-time tracing and monitoring of SARS-CoV-2 transmission through the large-scale global phylogenetic analysis of the ever-increasing SARS-CoV-2 genome sequences.

## Materials and Methods

### Implementation of TIPars

After assigning the ancestral sequences at every internal node and taxa sequences at external nodes, TIPars inserts a set of new samples into the reference phylogenetic tree sequentially based on parsimony criteria.

For a query sequence Q, TIPars computes the minimal substitution score against every branch in the tree. While inserting query Q into to the branch A-B (parent node - child node) at a potential newly added node P (Fig. 1A), the minimal substitution score is the sum of substitution scores that sequence Q differs from both sequence A and sequence B based on a specific substitution scoring table based on the IUPAC nucleotide ambiguity codes (table S3). The single branch with the minimum substitution score *σ* is reported as the best placement.

However, in terms of multiple placements where more than one branch have the same minimum substitution score, TIPars applies simple but practical rules to filter them to a single best placement such that multiple queries would be inserted sequentially based on one resulting tree. The first priority is to select the branch with node A containing the most numbers of child nodes. The second priority is to select the branch with node A of the lowest node height, that is the total branch length on the longest path from the node to a leaf (Suchard et al., 2018). Finally, in the case where the ambiguity cannot be resolved by the first two priorities, TIPars just turns to a pick up randomly. Even though TIPars will filter out multiple placements, these potential placements will also be printed out for user notice.

We proposed a local estimation model to calculate the pendant length of the newly introduced branch P-Q (*l_P-Q_*) which is considering the branch lengths of the local triplet subtree (A,(B,Q)) (Fig. 1A). Pendant length is defined as *l_P-Q_* = *σ*/(*δ_A_* + *δ_B_*)\**l_A-B_*, where *δ_A_* and *δ_B_* are the unique mismatch substitution scores of Q to A and B, and *l_A-B_* is the original length of branch A-B. The location of P on branch A-B is determined by the ration of *δ_A_* and *δ_B_*, i.e., Distal length: *l_A-P_* = *δ_A_*/(*δ_A_* + *δ_B_*)\**l_A-B_*, and Sibling length: *l_B-P_* = *δ_B_*/(*δ_A_* + *δ_B_*)\**l_A-B_*. The ancestral sequence of node P is estimated by majority vote of the nucleotide bases of sequence A, B and Q. To retain the topology of reference tree, a potential nucleotide base of Q will be only derived from A or B. For a special case of *l_A-B_* is zero but *σ* is not, TIPars will consider upper branch of A’s parent to A for scaling.

We implemented TIPars using Java with BEAST library (Suchard et al., 2018). Both FASTA and VCF formats are acceptable for loading sequences while NEWICK format is for the tree file. FASTA file is the default setting, but VCF file is more memory efficient for large dataset of high similar sequences, e.g. SARS-CoV-2 virus. To convert a FASTA file to VCF file with all sequence mutations, i.e. insertion, deletion and substitution, we used a Python package PoMo/FastaToVCF.py (Schrempf, Minh, De Maio, von Haeseler, & Kosiol, 2016).

### Benchmark datasets preparation

Unmasked SARS-CoV-2 MSA from GISAID was downloaded on 6th July 2021. Then all SARS-CoV-2 viral genome sequences collected before 1st January 2021 were extracted from the MSA. In order to ensure the sequences used for downstream analysis were complete, SARS-CoV-2 genomes with sequence length < 29,000 bp and > 0.5% Ns were removed (namely GISAID202101). To ensure that the global phylogenetic diversity is well represented in the sub-sampled dataset, sequences from all lineages as designated by the PANGO nomenclature system (Rambaut et al., 2020) were sub-sampled. Where fewer than 50 sequences of a given lineage were found in the global dataset, all sequences of the lineage were included. This resulted in a final sub-sampled dataset of 96,020 sequences from 1,249 PANGO lineages, with hCoV-19/Wuhan/WIV04/2019/EPI_ISL_402124 included as the reference genome (namely SARS2-100k). The SARS2-100k reference tree was then built using IQ-TREE2 with GTR model using the EPI_ISL_402124 as root. Ancestral sequences of each internal node were estimated using PastML with the MSA and the IQ-TREE2 generated tree as input.

Three small but representative nucleotide sequence datasets namely, bacterial 16S rRNA (16S), hemagglutinin genes of human seasonal influenza A viruses (H3N2), and Newcastle disease virus genomes (NDV), were prepared for programs performance comparison. The 16S dataset was downloaded from Genomes OnLine Database (Mukherjee et al., 2019) and randomly down-sampled to 800 sequences. HA sequences of 800 H3N2 viruses were randomly extracted from Influenza Research Database (Zhang et al., 2017). The 235 NDV sequences were downloaded from GenBank. Alignments were constructed using MUSCLE (Edgar, 2004). Reference trees of these datasets were built using RAxML (Stamatakis, 2014) standard hill-climbing heuristic search with 100 multiple inferences and GTRGAMMA model. Ancestral sequences were estimated using ML joint method (Pupko et al., 2000).

### Novel SARS-CoV-2 query sequence dataset

To generate novel query sequences for the 20 sets of 100, 1000, 5000 and 10000 sequences, SARS-CoV-2 genomes that were not included in the SARS2-100k dataset were randomly selected from the GISAID202101 dataset. Selected sequences were then aligned to the SARS2-100k sequences alignment by opening necessary gaps to obtain the full-length MSA. The newly selected sequences were extracted to obtain the final query sample sets. Corresponding new gaps were also added back to the ancestral sequence alignment for each dataset generated. PANGO lineages data for the novel SARS-CoV-2 query sequences and the taxa of reference tree was downloaded from GISAID on 6th July 2021.

### Benchmark programs

We compared TIPars to four state-of-the-art phylogenetic placement tools, namely UShER, EPAng, IQ-TREE2 and PAGAN2 while EPA-ng only works for single taxon placement and IQ-TREE2 and PAGAN2 were only used for multiple taxa insertion.

For the SARS2-100k dataset, only TIPars and UShER were considered as the other programs were not able to complete the computation within a reasonable runtime (Table 1). For the three smaller datasets, we compared all of them comprehensively. Details of the commands used for different programs could be found in table S8.

TIPars, UShER and EPA-ng would report multiple placements for single taxon insertion. The marked best placements of TIPars and UShER by themselves were used for our accurac evaluation. EPA-ng reports its results sorted by log-likelihood, so the placement with the highest log-likelihood was applied for assessment.

For any tools that accept only binary tree, i.e., EPA-ng and PAGAN2, we first converted the original polytomous tree to a binary tree using the Ape R package (Paradis & Schliep, 2019).

When adding unaligned query samples, it is suggested to align them to the MSA of taxa and ancestral sequences in the reference tree using MAFFT (‘--add’ option) (Katoh & Standley, 2013).

### Evaluation metrics

For single taxon placement evaluation, we first pruned one taxon from the reference tree and re-inserted it back. To assess the consistency between placement algorithms and the typical tree-constructing approach, we proposed using Robinson–Foulds (RF) Distance as a measure of the tree topology accuracy, as calculated by TreeCmp (Bogdanowicz, Giaro, & Wróbel, 2012). When the RF distance between a hypothetical tree and the reference tree is zero, the topology of the hypothetical tree is the same as the reference tree which means the algorithm inserts the query sample into the reference tree topological correctly. Another performance comparison with different true positive definition was conducted for binary trees derived from trees with polytomy using the measurement of whether sister node sets are identical to reference (Y. Turakhia et al., 2020).

For multiple taxa insertion evaluation, we randomly pruned a set of taxa from the reference tree and re-inserted them back. In addition to using RF distance to compare the hypothetical tree against the reference tree, we also calculated the log-likelihood of the hypothetical tree as a measurement of the accuracy of the taxa insertions. We applied two methods to compute log-likelihoods including FastTree2 (double-precision version) (Gamma20 Log-Likelihood) (Price et al., 2010) for optimized branch length, and IQ-TREE2 (Log-Likelihood (Fixed Br)) for fixed branch length.

EPA-ng outputs the placement information (placed branch, distal length, and pendant length) for a query without the construction of the final tree. In order to compute the RF distance, we assisted EPA-ng in inserting the query into the reference tree to generate the hypothetical tree.

IQ-TREE2 and PAGAN2 support initial tree, but they are not exactly based on the input tree topology for construction, so RF distance to original reference tree is not suitable for them. Note that UShER outputs the final constructed tree using the number of mutations as branch length (otherwise no branch length would be specified at branches modified), so we modified its branch length as number of mutations divided by alignment length in calculation of log-likelihood with fixed branch length model.

### Statistics

99% t-test confident intervals and 99% paired t-test p-value (right tail) for the results of TIPars against other programs were computed by Matlab R2013b. All violin graphs were generated by R 4.1.1 using the package *ggstatsplot* (Patil, 2021). Illustration and annotation of phylogenetic trees were done using the R package *ggtree* (G. Yu, Smith, Zhu, Guan, & Lam, 2017).

## Data and materials availability

SARS2-CoV-2 data used in this work were all downloaded from GISAID (https://www.gisaid.org/). TIPars is available at https://github.com/id-bioinfo/TIPars.

## Acknowledgments

We gratefully acknowledge the following Authors from the Originating laboratories responsible for obtaining the specimens and the Submitting laboratories where genetic sequence data were generated and shared via GISAID Initiative, on which this research is based. A full acknowledgement table can be found with two EPI_SET-IDs, i.e., EPI_SET_20211201vz and EPI_SET_20211206tc, in Data Acknowledgement Locator under GISAID resources (https://www.gisaid.org/).

This project is supported by the Hong Kong Research Grants Council’s General Research Fund (17150816), the National Natural Science Foundation of China’s Excellent Young Scientists Fund (Hong Kong and Macau) (31922087), the Health and Medical Research Fund (COVID1903011-WP1), the Innovation and Technology Commission’s InnoHK funding (D^2^4H), and the Guangdong Government for the funding supports (2019B121205009, HZQB-KCZYZ-2021014, 200109155890863, 190830095586328 and 190824215544727).

