## Supplementary Materials for "Robust expansion of phylogeny for fast-growing genome sequence data"

**Supplementary materials include:**

Fig. S1 to S4

Table S1 to S8

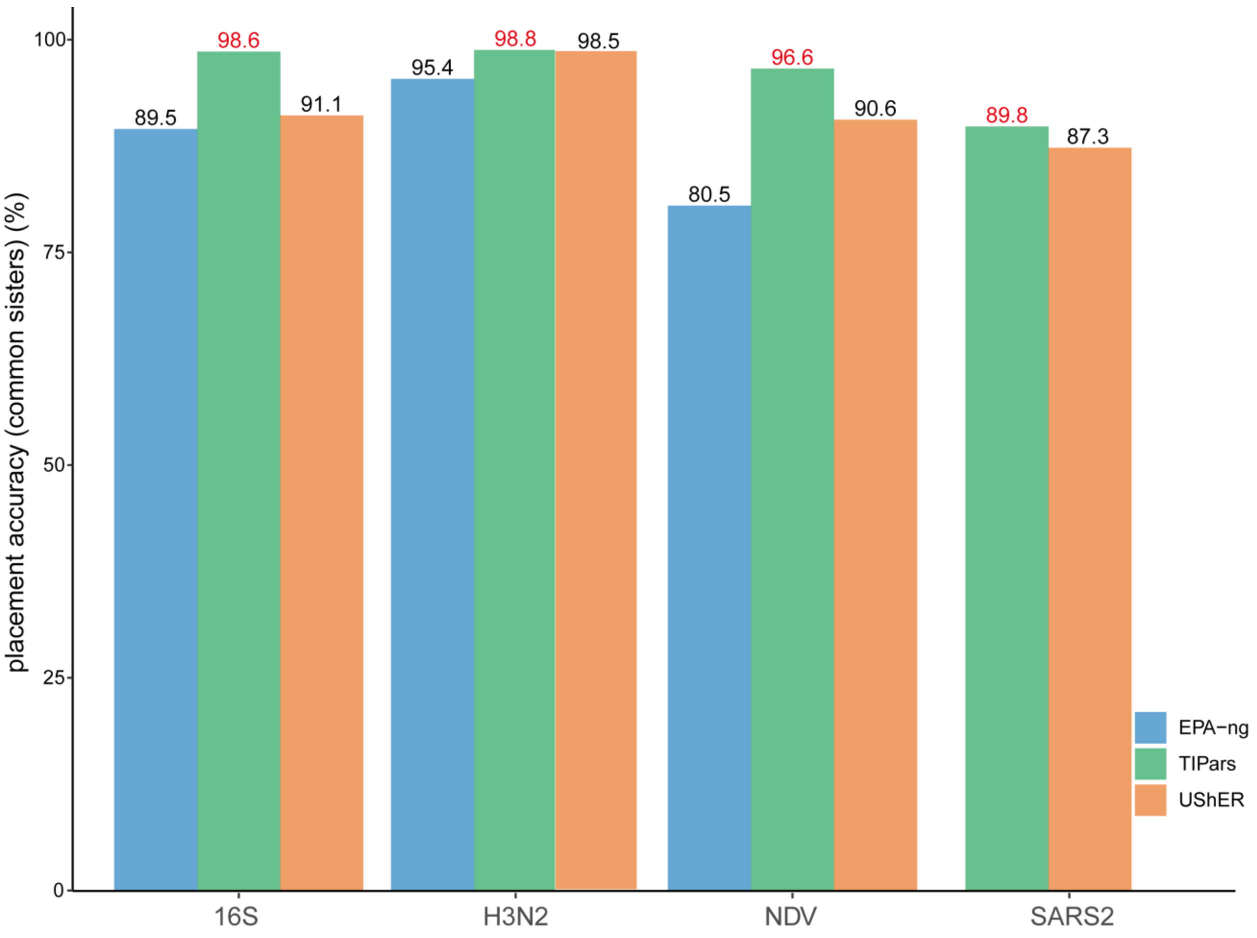


**Fig. S1. Single taxon placement performance measured by common-sisters.** Bar charts represent the accuracy of single taxon placement on 16S, H3N2, NDV and SARS2-100k datasets for TIPars, UShER and EPA-ng respectively. Accuracy is indicated on top of each bar and the highest accuracy in each dataset is highlighted in red. A true positive of common sisters is defined as the query has sister nodes in the resulting tree that are a subset of sister nodes of this query before removed from the reference tree.

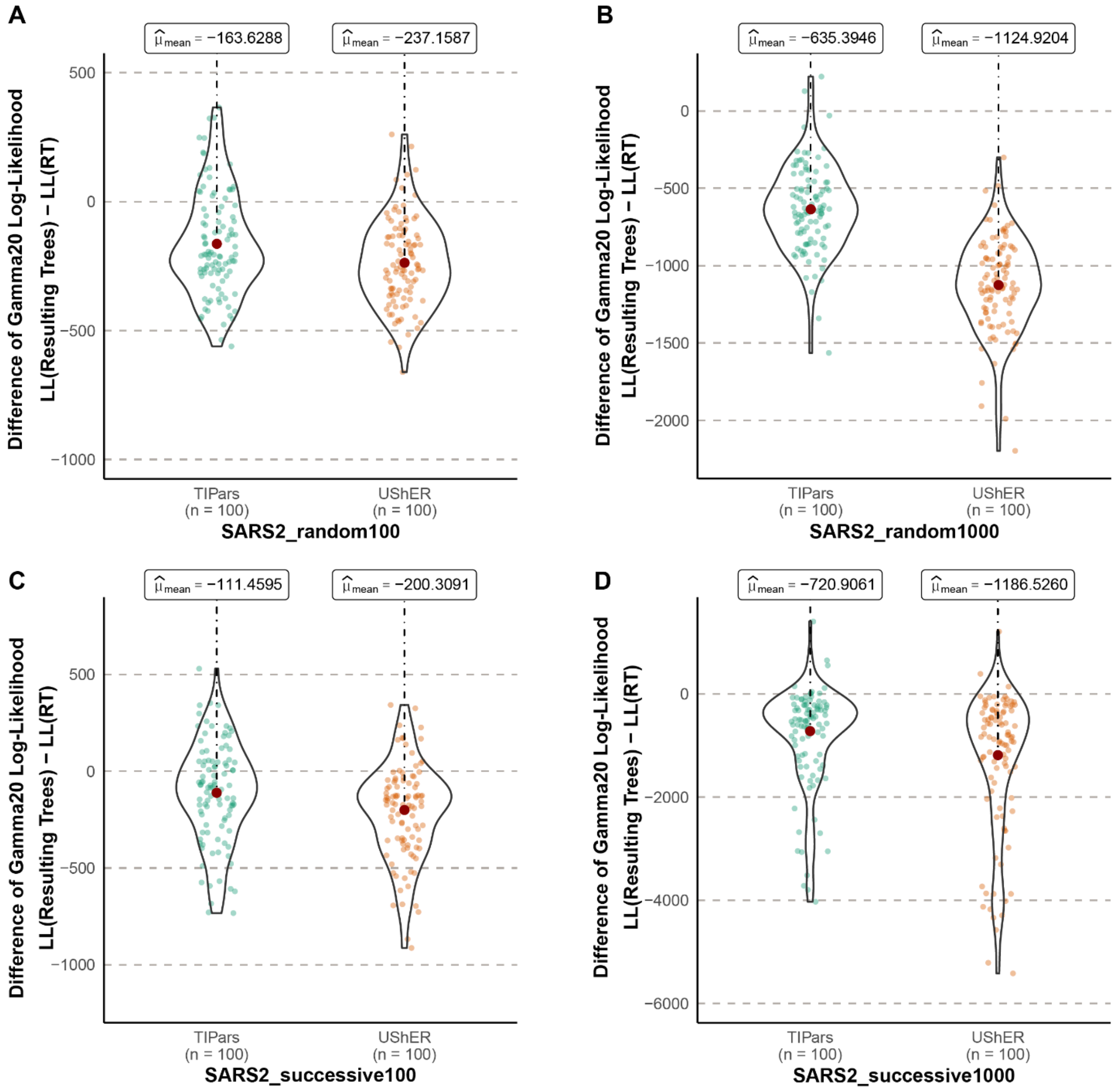


**Fig. S2. Paired differences of the Gamma20 log-likelihood (LL) between the optimized resulting trees and reference tree (RT) on SARS2-100k dataset.** Violin graphs show the distribution of paired differences for the random 100 (**A**), 1000 (**B**) and successive 100 (**C**), 1000 (**D**) multiple sequences insertions.

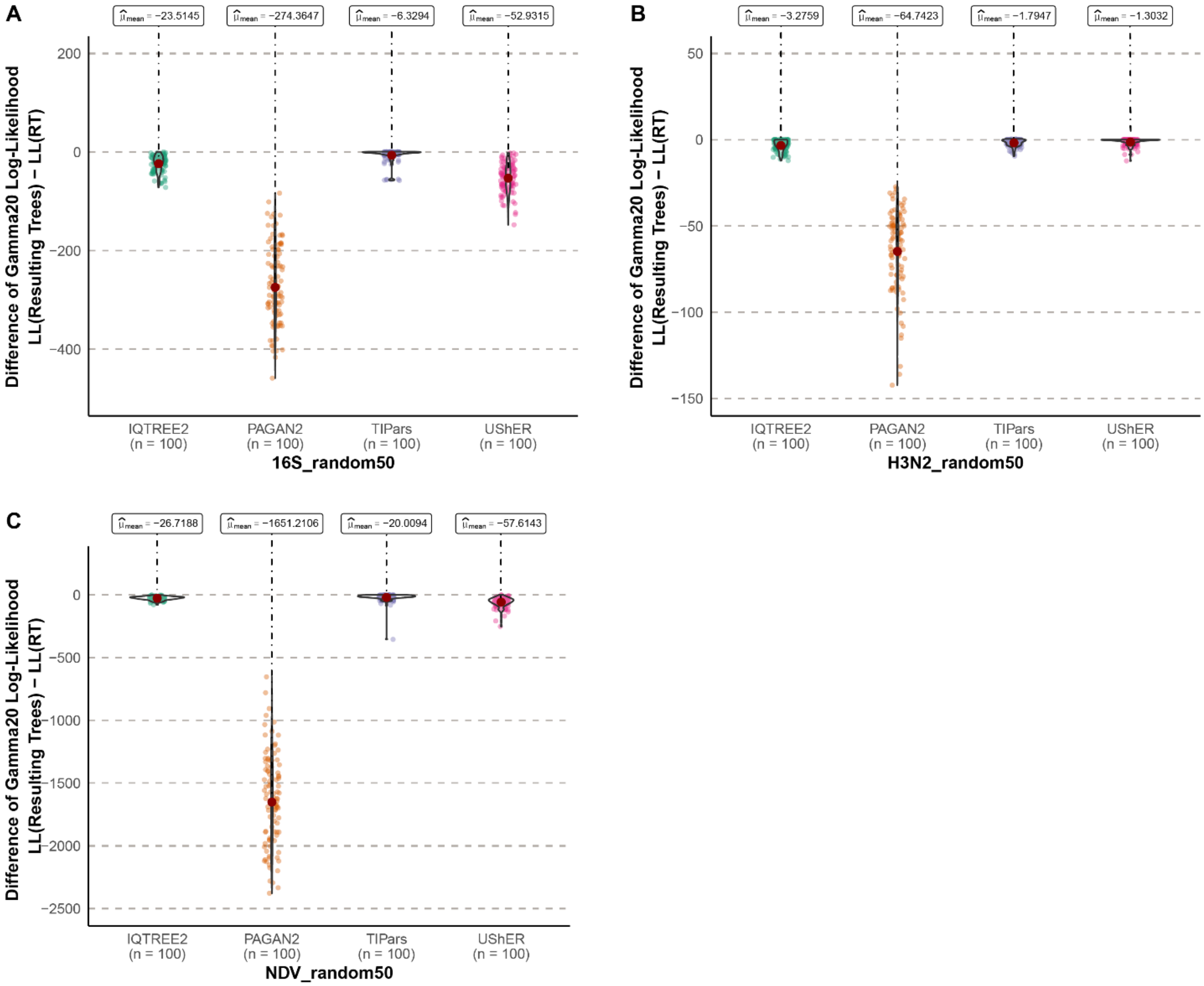


**Fig. S3. Paired differences of the Gamma20 log-likelihood (LL) between the optimized resulting trees and reference tree (RT).** Violin graphs show distribution of the paired differences for random 50 multiple sequences insertions on 16S (**A**), H3N2 (**B**) and NDV (**C**).

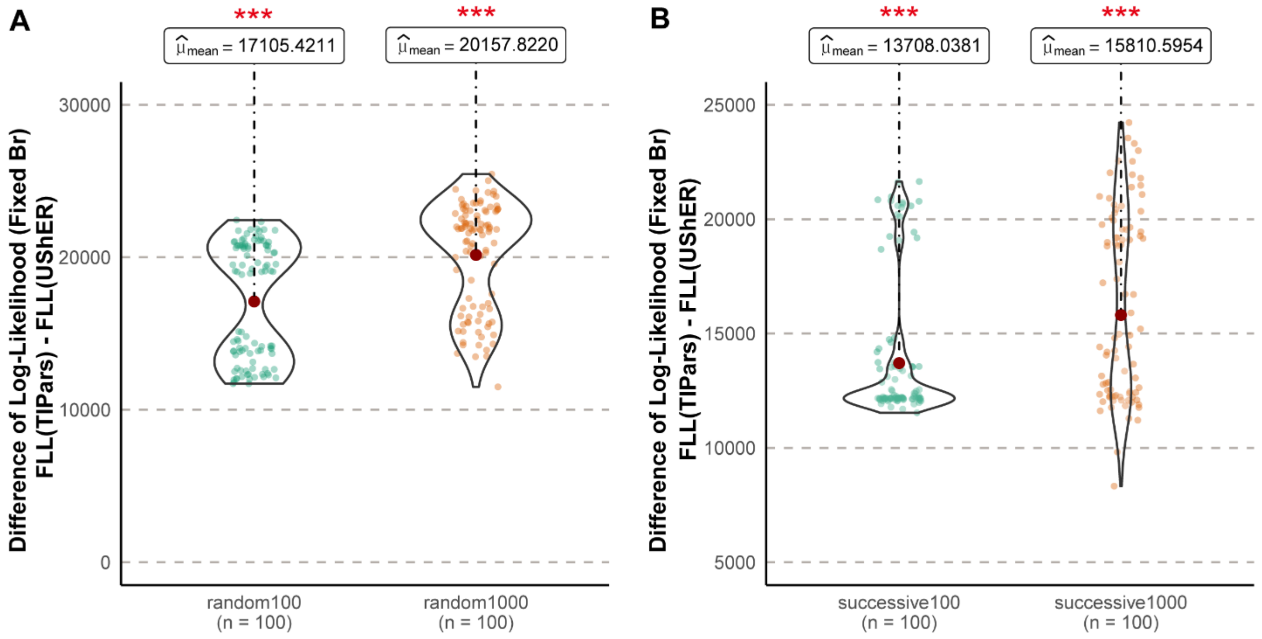


**Fig. S4. Paired differences of the log-likelihood (FLL) between the resulting trees with fixed branch length generated by TIPars and UShER.** Violin graphs show the distribution of paired differences for the random 100, 1000 (**A**) and successive 100, 1000 (**B**) multiple sequences insertions on SARS2-100k dataset. P-value for the right-sided paired t-test is indicated by the asterisk on top of each violin diagram, where $p<0.001$ is indicated by three red asterisks (***).

**Table. S1. Average running time and memory used through 10 repeated runs of inserting/placing 50 16S samples onto 16S 800-taxa reference tree.** Tests were running in a server of 64 Intel Xeon Gold 6242 CPU cores and 1500 GB RAM. We also compared TIPars with UShER in a general computer of 8 CPU cores. TIPars ran with a JAVA setting of -Xmx1G loaded with FASTA files. The running time of UShER contains its necessary computation of 'mutation-annotated tree'. EPA-ng insertion time is not available because of its limitation to the precision of second. N/A indicates that data are not applicable.

| **Tools** | **CPU cores assigned** | **Mean insertion time (Seconds)** | **Mean running time (Seconds)** | **Mean peak memory (GB)** |
|---|---|---|---|---|
| **TIPars** | 64 | 0.83 | 1.22 | 0.11 |
| **TIPars** | 8 | 0.36 | 0.69 | 0.07 |
| **UShER** | 64 | 0.47 | 0.65 | 0.02 |
| **UShER** | 8 | 1.87 | 2.08 | 0.01 |
| **EPA-ng** | 64 | N/A | 0.69 | 0.78 |
| **IQ-TREE2** | 64 | N/A | 2.11 | 0.05 |
| **PAGAN2** | 64 | N/A | 40.54 | 0.50 |

**Table. S2. Average pairwise genetic distance per site for each benchmark dataset**. A random 10% sub-samples of SARS-CoV-2 datasets were used to compute the genetic distance, while all sequences in other datasets were used for genetic distance calculation.

| Dataset | Number of sequences | Alignment length (bp) | Average pairwise genetic distance (substitutions per site) |
|---|---|---|---|
| **SARS2-660k** | 659,885 | 29,891 | 1.26E-03 |
| **SARS2-100k** | 96,020 | 30,545 | 7.57E-04 |
| **16S** | 800 | 1,432 | 2.09E-01 |
| **H3N2** | 800 | 1,698 | 3.30E-02 |
| **NDV** | 235 | 15,072 | 1.22E-01 |

**Table S3. IUPAC code substitution table**

| | A | C | G | T | R | Y | M | K | S | W | H | B | V | D | N | - |
|---|---|---|---|---|---|---|---|---|---|---|---|---|---|---|---|---|
| **A** | 0 | 1 | 1 | 1 | 0 | 1 | 0 | 1 | 1 | 0 | 0 | 1 | 0 | 0 | 0 | 0 |
| **C** | 1 | 0 | 1 | 1 | 1 | 0 | 0 | 1 | 0 | 1 | 0 | 0 | 0 | 1 | 0 | 0 |
| **G** | 1 | 1 | 0 | 1 | 0 | 1 | 1 | 0 | 0 | 1 | 1 | 0 | 0 | 0 | 0 | 0 |
| **T** | 1 | 1 | 1 | 0 | 1 | 0 | 1 | 0 | 1 | 0 | 0 | 0 | 1 | 0 | 0 | 0 |
| **R** | 0 | 1 | 0 | 1 | 0 | 1 | 0 | 0 | 0 | 0 | 0 | 0 | 0 | 0 | 0 | 0 |
| **Y** | 1 | 0 | 1 | 0 | 1 | 0 | 0 | 0 | 0 | 0 | 0 | 0 | 0 | 0 | 0 | 0 |
| **M** | 0 | 0 | 1 | 1 | 0 | 0 | 0 | 1 | 0 | 0 | 0 | 0 | 0 | 0 | 0 | 0 |
| **K** | 1 | 1 | 0 | 0 | 0 | 0 | 1 | 0 | 0 | 0 | 0 | 0 | 0 | 0 | 0 | 0 |
| **S** | 1 | 0 | 0 | 1 | 0 | 0 | 0 | 0 | 0 | 1 | 0 | 0 | 0 | 0 | 0 | 0 |
| **W** | 0 | 1 | 1 | 0 | 0 | 0 | 0 | 0 | 1 | 0 | 0 | 0 | 0 | 0 | 0 | 0 |
| **H** | 0 | 0 | 1 | 0 | 0 | 0 | 0 | 0 | 0 | 0 | 0 | 0 | 0 | 0 | 0 | 0 |
| **B** | 1 | 0 | 0 | 0 | 0 | 0 | 0 | 0 | 0 | 0 | 0 | 0 | 0 | 0 | 0 | 0 |
| **V** | 0 | 0 | 0 | 1 | 0 | 0 | 0 | 0 | 0 | 0 | 0 | 0 | 0 | 0 | 0 | 0 |
| **D** | 0 | 1 | 0 | 0 | 0 | 0 | 0 | 0 | 0 | 0 | 0 | 0 | 0 | 0 | 0 | 0 |
| **N** | 0 | 0 | 0 | 0 | 0 | 0 | 0 | 0 | 0 | 0 | 0 | 0 | 0 | 0 | 0 | 0 |
| **-** | 0 | 0 | 0 | 0 | 0 | 0 | 0 | 0 | 0 | 0 | 0 | 0 | 0 | 0 | 0 | 0 |

**Table S4. RF distance per query taxon in every testing set.** For each taxon in every set of successively removed taxa (as query samples), we pruned other queries except itself from the resulting tree and computed the RF distance to the corresponding reference tree. The RF distance per query taxon for statistics is the mean of those of all queries in each testing set.

| | successive100 | | successive1000 | |
|---|---|---|---|---|
| | **TIPars** | **UShER** | **TIPars** | **UShER** |
| **Mean** | 4.81 | 4.14 | 6.51 | 6.07 |
| **Confidence** | 3.07 | 2.56 | 4.72 | 4.47 |
| **interval** | 6.54 | 5.73 | 8.31 | 7.67 |
| **P-value** | N/A | 0.12 | N/A | 0.21 |

**Table. S5. Running time and memory used for optimizing branch length of a SARS-CoV-2 100k taxa tree by FastTree2 (double-precision version) for 100 times.** The computation was conducted in a server of 8 Intel Xeon Gold 6242R CPU cores.

| Mean running time (HH:MM:SS) | Mean peak memory (GB) | Time range over 100 runs | Memory range over 100 runs (GB) |
|---|---|---|---|
| 10:54:28 | 126 | 10:18:48 - 12:34:04 | 125 - 126 |

**Table S6. PANGO lineages accuracy of 20 sets of 100, 1000, 5000 and 10000 novel samples insertion to the SARS-CoV-2 100k taxa tree.**

| | 100 | | 1000 | | 5000 | | 10000 | |
|---|---|---|---|---|---|---|---|---|
| **Set_Id** | **TIPars** | **UShER** | **TIPars** | **UShER** | **TIPars** | **UShER** | **TIPars** | **UShER** |
| **1** | 96.91 | 96.91 | 90.96 | 90.26 | 91.95 | 91.57 | 91.14 | 90.77 |
| **2** | 89.90 | 89.90 | 91.34 | 90.33 | 91.52 | 91.19 | 91.36 | 91.24 |
| **3** | 92.71 | 92.71 | 89.95 | 90.05 | 92.00 | 91.78 | 91.38 | 91.17 |
| **4** | 95.92 | 95.92 | 92.08 | 91.88 | 91.65 | 90.98 | 91.29 | 91.07 |
| **5** | 93.94 | 91.92 | 92.07 | 90.76 | 91.91 | 91.61 | 91.60 | 91.36 |
| **6** | 92.93 | 93.94 | 91.99 | 91.58 | 91.14 | 91.08 | 91.05 | 90.79 |
| **7** | 87.91 | 87.91 | 92.31 | 91.60 | 91.72 | 91.31 | 91.10 | 91.06 |
| **8** | 87.63 | 86.60 | 92.67 | 92.37 | 91.90 | 91.58 | 91.04 | 90.76 |
| **9** | 92.93 | 92.93 | 90.86 | 90.05 | 91.82 | 91.58 | 91.80 | 91.55 |
| **10** | 86.46 | 85.42 | 92.00 | 91.29 | 91.73 | 91.12 | 92.01 | 91.57 |
| **11** | 89.90 | 88.89 | 91.77 | 91.37 | 91.61 | 91.30 | 91.63 | 91.34 |
| **12** | 90.00 | 91.00 | 91.13 | 90.93 | 91.70 | 91.60 | 91.86 | 91.47 |
| **13** | 87.37 | 87.37 | 91.23 | 90.32 | 90.64 | 90.30 | 91.40 | 90.96 |
| **14** | 91.84 | 89.80 | 92.93 | 92.73 | 91.04 | 90.67 | 91.60 | 91.15 |
| **15** | 88.00 | 86.00 | 92.50 | 91.89 | 91.43 | 90.96 | 91.29 | 91.10 |
| **16** | 92.00 | 92.00 | 90.17 | 89.46 | 91.37 | 91.00 | 91.38 | 91.04 |
| **17** | 89.58 | 89.58 | 91.51 | 91.20 | 91.11 | 90.86 | 91.17 | 91.32 |
| **18** | 96.94 | 96.94 | 93.80 | 93.19 | 91.82 | 91.47 | 91.41 | 91.13 |
| **19** | 97.00 | 97.00 | 90.51 | 90.10 | 91.05 | 90.89 | 91.49 | 91.33 |
| **20** | 95.00 | 95.00 | 92.65 | 92.25 | 91.54 | 91.12 | 91.73 | 91.34 |

**Table S7. Cases of Gamma20 Log-Likelihoods with equal RF distance between TIPars and UShER.** There is no case in random1000 dataset.

| | | RF distance | | Gamma20 Log-Likelihhod | | |
|---|---|---|---|---|---|---|
| **Dataset** | **Index** | **TIPars** | **UShER** | **TIPars** | **UShER** | **Difference** |
| **random100** | 28 | 102.5 | 102.5 | -1953459.706 | -1953464.289 | 4.583 |
| **random1000** | NONE | N/A | N/A | N/A | N/A | N/A |
| **successive100** | 3 | 4 | 4 | -1953115.096 | -1953451.73 | 336.634 |
| | 13 | 9.5 | 9.5 | -1953140.877 | -1953361.722 | 220.845 |
| | 44 | 0 | 0 | -1952829.626 | -1953129.241 | 299.615 |
| | 66 | 14 | 14 | -1953013.427 | -1953142.914 | 129.487 |
| | 98 | 1 | 1 | -1953111.782 | -1952933.306 | -178.476 |
| **successive1000** | 11 | 121.5 | 121.5 | -1953766.679 | -1955187.046 | 1420.367 |
| **16S** | 31 | 7.5 | 7.5 | -165389.527 | -165466.808 | 77.281 |
| **H3N2** | 1 | 1.5 | 1.5 | -23533.245 | -23533.245 | 0 |
| | 5 | 0 | 0 | -23532.111 | -23532.111 | 0 |
| | 8 | 0 | 0 | -23532.111 | -23532.111 | 0 |
| | 15 | 0.5 | 0.5 | -23532.111 | -23532.111 | 0 |
| | 17 | 0.5 | 0.5 | -23532.111 | -23532.111 | 0 |
| | 20 | 2 | 2 | -23534.878 | -23536.814 | 1.936 |
| | 21 | 0.5 | 0.5 | -23532.111 | -23532.114 | 0.003 |
| | 32 | 1 | 1 | -23532.111 | -23532.111 | 0 |
| | 37 | 0.5 | 0.5 | -23532.111 | -23532.111 | 0 |
| | 44 | 0.5 | 0.5 | -23534.878 | -23536.298 | 1.42 |
| | 48 | 0 | 0 | -23532.111 | -23532.111 | 0 |
| | 49 | 0 | 0 | -23532.111 | -23532.111 | 0 |
| | 52 | 2 | 2 | -23533.366 | -23533.363 | -0.003 |
| | 53 | 0.5 | 0.5 | -23532.114 | -23532.111 | -0.003 |
| | 59 | 3 | 3 | -23534.378 | -23535.385 | 1.007 |
| | 69 | 0 | 0 | -23532.105 | -23532.111 | 0.006 |
| | 77 | 4.5 | 4.5 | -23533.181 | -23533.181 | 0 |
| | 82 | 0 | 0 | -23532.111 | -23532.111 | 0 |
| **NDV** | 6 | 9 | 9 | -274517.545 | -274606.407 | 88.862 |
| | 29 | 7 | 7 | -274502.29 | -274502.95 | 0.66 |

**Table S8. Running command of different tested tools**

**Tree Placement/Insertion**

| | |
|---|---|
| TIPars | `$tipars -t $treeFile -s $taxaFile -a $ancFile -q $queryFile -o $outFile` |
| UShER | `$usher --vcf $taxa&queryFile --tree $treeFile --write-uncondensed-final-tree --outdir $outdir` |
| IQ-TREE2 | `$iqtree -g $treeFile -s $taxa&queryFile -n 0 -m GTR -fixbr -pre iqtree2` |
| EPA-ng | `$epang --ref-msa $taxaFile --tree $treeFile --query $queryFile --model GTR -T 0` |
| PAGAN2 | `$pagan --ref-treefile $treeFile --ref-seqfile $taxaFile --queryfile $queryFile --guidetree --one-placement-only` |

**Log-Likelihood**

FastTree2 (double-precision version)

```
$fasttree -gamma -nt -gtr -nome -mllen -intree $treeFile -log $logFile
$taxaFile > $opt_treeFile
```

IQ-TREE2 (fixed branch length setting)

```
$iqtree -s $taxaFile -keep-ident -n 0 -m GTR -fast -nt 64 -blmin 0.0000000001
-me 0.05 --suppress-list-of-sequences --suppress-duplicate-sequence --no-opt-
gamma-inv -pre iqtree2 -fixbr -z $treeFiles
```

**RF distance**

TreeCmp

```
java -jar  $treecmp.jar -d rf -i $testTreeFile -r $refTreeFile -o $outFile
```
